# Changes in nitrogen pools in the maize-soil system after urea or straw application to a typical intensive agricultural soil: A ^15^N tracer study

**DOI:** 10.1101/2020.11.09.373928

**Authors:** Jie Zhang, Ping He, Dan Wei, Liang Jin, Lijuan Zhang, Ling Li, Shicheng Zhao, Xinpeng Xu, Wei Zhou, Peter Christie, Shaojun Qiu

## Abstract

A ^15^N maize pot experiment was conducted to compare the N value of fertilizer alone and fertilizer combined with straw at an equivalent N rate. The four treatments were control (CK), ^15^N-urea, ^15^N-urea plus straw, and ^15^N-straw plus urea. Soil N pools, maize N and their ^15^N abundance were determined during maize growth. At maturity 26.0% of straw N was assimilated by maize in the urea plus straw treatment. From the eighth leaf stage to maturity, urea plus straw had a significantly (P < 0.05) higher concentration and percentage of exogenous substrate N present as soil total N (TN), particulate organic N (PON), and mineral associated total N (MTN) in bulk and rhizosphere soils than the urea-only treatment. From silking to maturity in the urea plus straw treatment, rhizosphere soil significantly (P < 0.05) increased the percentage of exogenous substrate N present as inorganic N (Inorg-N) and MTN, and significantly (P < 0.05) decreased that present as PON and microbial biomass N (MBN) compared with the bulk soil. From the eighth leaf stage to maturity, rhizosphere soil significantly (P < 0.05) increased the percentage of straw N present as Inorg-N and MTN except for MTN at the silking stage, and significantly decreased (P < 0.05) that present as PON compared with the bulk soil. Overall, straw was an available N source to the crop, and the increase in straw N availability needs to be considered from the interaction of fertilization practices and the crop rhizosphere.

## 1. Introduction

Fertilizer nitrogen (N) has made a very large contribution to crop production to meet the food demand of an increasing world population since World War II [1]. However, numerous environmental issues have emerged with increasing application rate of chemical fertilizer N, and long-term straw removal from agricultural land additionally aggravated the loss of inorganic nutrients and soil acidification, and straw burning leads to atmospheric pollution [2, 3]. Conversely, straw is an important source of carbon and the combined application of chemical fertilizer N with straw is an effective practice for the maintenance of soil fertility and promotion of the sustainable utilization of agricultural soils [4, 5]. The effects of straw C on soil fertility have been of greater interest than the availability of straw N for crop uptake. Consequently, the proportion of straw N that can be assimilated during crop growth under the combined application of chemical fertilizer and straw remains unclear.

The dynamics of crop N assimilation are closely linked with soil N supply and N flow from the bulk soil to the rhizosphere soil. In general, rhizosphere soil has a greater tendency to show N deficiency than bulk soil during crop growth, and then a N flow gradient forms from bulk soil to rhizosphere soil. However, the N flow rate changes with root growth and is regulated by living roots, root activity, soil nutrient transformations, and nutrient transfer from soil to crop [6, 7]. The chemical fertilizer N in the soil solution can be transferred from the bulk soil to the rhizosphere soil and is directly assimilated by crops, but straw N assimilation by crops occurs only after decomposition of the straw by microorganisms. Microbial mineralization of straw usually requires additional chemical fertilizer N application due to the high C:N ratio in straw but there is little information on whether the mineralization of straw N in the rhizosphere soil increases or decreases compared to the bulk soil because root exudates can promote microbial N immobilization and also substantial crop N assimilation generates relative N deficiency during crop growth. It is therefore necessary to compare the behaviors of both straw N and chemical fertilizer N in the rhizosphere soil and the bulk soil during crop growth.

Non-leguminous crops assimilate N mainly from the soil and soil N can be differentiated into different labile pools. In agricultural ecosystems, soil inorganic N (Inorg-N) is derived mainly from chemical fertilizers and can be immobilized in or released from soil organic N pools [8, 9]. The labile organic N pools are soil microbial biomass N (MBN), dissolved organic N (DON), and particulate organic N (PON) [9]. MBN regulates the immobilization of Inorg-N and the mineralization of soil organic N. Inorg-N and DON can be directly or indirectly assimilated by crops and microorganisms, and DON is derived from microbial metabolism and root exudation and further decomposes to Inorg-N via microbial activity [10–12]. PON represents partly decomposed plant residues and is an N source for soil microorganisms [13, 14]. When PON is removed from the whole soil the residual N is regarded as mineral associated total N (MTN). MTN holds the largest proportion of the soil total N [15] and retains N substrates by sorption on mineral surfaces or the formation of organic-mineral complexes [16, 17]. These N pools can therefore reflect soil biological, chemical, and physical properties.

In agricultural systems chemical fertilizer N application can stimulate soil organic matter mineralization and promote soil N transformation among the different soil N pools. In addition, straw returned to the soil acts as “temporal POM” [18, 19], provides a C source for microorganisms, and further drives soil N transformations [20, 21]. When straw and chemical fertilizer N are combined, the microbial immobilization of chemical fertilizer N is induced by straw C and interactions between straw N and chemical fertilizer N can decrease the loss of fertilizer N and promote the transformation of fertilizer N into soil N pools. However, chemical fertilizer N stimulates the microbial mineralization of straw N after combination with straw and the bioavailability of straw N increases [5]. In addition, competition for N occurs between microorganisms and crops accompanied by crop growth and this further alters N retention in different soil N pools. Overall, it is necessary to explore how different soil N pools change under the interactions among chemical fertilizer applications, straw applications, and crop growth.

The main objectives of the present study were to quantify the concentrations and percentages of chemical fertilizer N and straw N present in different soil N pools (Inorg-N, MBN, DON, PON, MTN), N uptake by different plant parts (shoots, grains, roots), the recoveries of chemical fertilizer N and straw N in different soil N pools and plant parts in mature maize plants, and to compare the differences in soil N pools between rhizosphere soil and bulk soil during maize growth after the application of fertilizer N and straw. A pot experiment was conducted with ^15^N-labeled urea and ^15^N-labeled maize straw in order to address these issues.

## 2. Materials and methods

### 2.1. Soil

Soil samples were collected from an arable field (0-20 cm depth) at Heilongjiang Academy of Agricultural Sciences near Harbin city, Heilongjiang province, northeast China (45°50’ N, 126°51’ E). The region is characterized by low temperatures and a continental monsoon climate with an average annual precipitation of 486 mm and mean annual temperature of 3.6 °C. The soil is classified as a Mollisol according to the USDA classification [22]. Selected soil physicochemical properties are shown in Table 1.

**Table 1.** Selected physicochemical properties of the chernozem soil studied.

| Organic C | Total N | Inorganic N | Available P | Available K | pH | Soil texture (%) |  |  |
| --- | --- | --- | --- | --- | --- | --- | --- | --- |
| (g kg<sup>-1</sup>) | (g kg<sup>-1</sup>) | (mg kg<sup>-1</sup>) | (mg kg<sup>-1</sup>) | (mg kg<sup>-1</sup>) | (units) | Sand | Silt | Clay |
| 19.4 | 1.8 | 22.8 | 9.6 | 155.9 | 6.9 | 27.4 | 48 | 24.6 |

### 2.2. Preparation of labeled and unlabeled straw

^15^N-labeled straw was obtained by conducting a greenhouse pot experiment. As previously reported by Qiu *et al.* (2012) [5], labeled straw with a relatively high ^15^N abundance was produced by growing maize (cv. Zhengdan 958) in soil with 30.16 % ^15^N-labeled (NH_4_)_2_SO_4_, NaH_2_PO_4_ and KCl at rates of 150 mg N, 65.5 mg P and 124.5 mg K per kg soil. Nitrogen limitation was minimized by applying the topdressing fertilizer at the 8th leaf and tasseling growth stages of the labeled maize at rates of 1 g plant^−1^ and 2 g plant^−1^ (^15^NH_4_)_2_SO_4_ at 30.16% ^15^N abundance, respectively. Maize grain formation was prevented by covering the corn cobs with paper bags before the silking stage. The unlabeled maize was grown in a field near the greenhouse with the same soil type and maize cultivar. The sowing and harvest dates of the unlabeled maize were the same as those of the labeled maize. After the experimental materials were harvested the total C, N, P, and K concentrations, and the ^15^N abundance of the labeled straw were 435.5 ± 1.2, 9.6 ± 0.2, 1.4 ± 0.02, and 11.2 ± 0.03 g kg^−1^, 45.3 ± 1.2, and 15.24 ± 0.06%, and the corresponding values of the unlabeled straw were 449.7 ± 0.17, 11.3 ± 0.2, 1.1 ± 0.01, and 14.2 ± 1.2 g kg^−1^, 41.7 ± 0.6, and 0.37 ± 0.0001%.

### 2.3. Experimental design

An equivalent N amendment pot experiment with a complete randomized design was carried out in a net house with a glass roof. There were four treatments: (1) control without N addition (CK); (2) ^15^N-labeled urea (^15^NU); (3) ^15^N-labeled urea plus straw (^15^NU + S); (4) ^15^N-labeled straw plus urea (^15^NS + U), and the ^15^NU + S and ^15^NS + U treatments are defined as U + S treatment. In the U + S treatment, exogenous substrate N was applied as the sum of urea N and straw N and the ratio of urea N to straw N was 6:4 according to Zhu and Wen (1997) [23]. The total exogenous substrate N added in the different N treatments was 1.5g pot^−1^, and the total P and K rates added in all treatments were 0.655 and 1.245 g pot^−1^, respectively. Each pot (30 cm height × 25 cm diameter) contained 10.0 kg soil (oven-dried basis) which was thoroughly mixed with exogenous substrate before sowing. The synthetic fertilizers used were urea, superphosphate, and potassium chloride, and the straw powder was passed through a 0.25-mm sieve. Equivalent N, P, and K rates in each pot were ensured by including the N, P, and K contents in the maize straw. There were 12 replicates of each treatment to allow destructive sampling at the 8th leaf (V8), silking (R1), milk (R3), and physiological maturity (R6) growth stages, i.e. 35th, 70th, 90th, and 111th days after the maize was sown.

The maize cultivar was ‘Zhengdan 958’. Two maize seeds were sown in the center of each pot and thinned to one plant after emergence. In order to prevent root growth along the inner wall of the pot, three PVC tubes (diameter 13.8 cm, height 35 cm) were inserted vertically into the soil in each pot. The base of each PVC tube was about 5 cm above the base of the pot, and the PVC tubes were about 5 cm from the pot inner wall. Each PVC tube had three holes 8 cm from the base, and the holes and the PVC tubes were enclosed with 0.5-mm mesh to prevent soil entering the PVC tube so that water in the tube flowed rapidly into the soil. Distilled water was added daily to each pot from the PVC tubes to constant weight, then the top of each tube was sealed with a rubber stopper to prevent loss of water. During maize growth the soil water content was 60 % using the weight balance method on a daily basis. Before the maize was sown the soil moisture and the air-dried soil were balanced for three days.

At each stage the shoots (aboveground) and roots (belowground) were separated from the maize upper node brace root. During maize growth the “less green” bottom leaves were cut, oven-dried, stored, and then mixed into the same plant sample when the maize plant was sampled. The sampled shoots, roots, and grains were oven-dried at 60 °C and then weighted with a balance.

Rhizosphere soil was sampled as described by Peng *et al.* (2012) [24]. Briefly, the roots were removed from the pot and shaken to remove the loosely attached soil with roots, then the soil adhering to the root system was placed in a paper bag, vigorously shaken, and brushed to collect the closely adhering soil with roots. The soil adhering to the roots was regarded as rhizosphere soil, and the remaining soil in each pot was mixed thoroughly and regarded as bulk soil. Any visible roots in either soil fraction were removed.

### 2.4. Sample analysis

Shoot N, root N, grain N and soil total N (TN) were determined with an elemental analyzer (Macro cube, Elementar, Hanau, Germany). Plant samples were passed through a 0.25-mm sieve and soil samples through a 0.15-mm sieve. The ^15^N abundance of shoot N, root N, grain N, and soil TN was determined using an isotope ratio mass spectrometer (Finnigan MAT251, Thermo Fisher, Waltham, MA).

Soil inorganic N (Inorg-N) was extracted with 1M KCl solution (1:5, W / V) on a reciprocal shaker for 1 h and then determined with a continuous flow analyzer (FIAstar 5000, FOSS, Hillerød, Denmark). For inorganic ^15^N abundance the 10 ml KCl-extracted solution was reduced using Devarda’s alloy and distilled. The distillates from the KCl extracts were quantified by titration and then acidified and oven-dried at 60 °C for N isotope analysis by mass spectrometry (Finnigan MAT251, Thermo Fisher, Waltham, MA) according to Hauck *et al.* (1996) [25].

Soil microbial biomass N (MBN) was determined using the CHCl_3_ fumigation-K_2_SO_4_ method as described by Brookes *et al.* (1985) [26]. Briefly, fresh soil was fumigated with CHCl_3_ for 24 h at 25 °C and then the N in fumigated and unfumigated samples was extracted with 0.5 M K_2_SO_4_ (1:4 w / v, 0.5 h). 20 ml fumigated and unfumigated solution was analyzed using the Kjeldahl method. Soil microbial biomass N was calculated as the difference in N concentration between fumigated and unfumigated samples divided by a conversion coefficient of 0.45 (Jenkinson 1988) [27]. Soil dissolved organic N (DON) was calculated by subtracting Inorg-N from the N concentration in unfumigated solution extracted with 0.5 M K_2_SO_4_ [9, 28]. The ^15^N abundance of fumigated and unfumigated solution was determined as ^15^N abundance of Inorg-N as described above.

Particulate organic matter and mineral associated matter were fractionated as described by Bronson *et al.* (2004) [29]. Briefly, fresh soil at each maize stage, equivalent to 25 g oven-dried soil (< 2 mm), was dispersed in 5% sodium hexametaphosphate solution (soil:solution ratio 1:4, w / v) on a reciprocal shaker for 60 min. The slurry was sieved using a 53-μm mesh until the deionized water became clear. The < 53 and ≥ 53 μm soil fractions were transferred to beakers separately and oven-dried at 60 °C. Nitrogen in both isolated soil fractions is defined as particulate organic N (PON, > 53 μm) and mineral associated total N (MTN, < 53 μm) and their N concentrations were determined using an elemental analyzer after particulate organic matter and mineral associated matter were passed through a 0.15-mm sieve (Macrocube, Elementar, Hanau, Germany). The ^15^N abundance of PON and MTN was determined as total ^15^N as described above.

### 2.5. Calculations

The concentration of exogenous substrate N present as MBN was calculated by the difference between the fumigated N concentration and the unfumigated N concentration derived from urea or straw, and then divided by a conversion coefficient of 0.45. Correspondingly, DON derived from urea or straw was calculated by the difference between unfumigated N content derived from urea or straw and Inorg-N derived from urea or straw.

The concentration and percentage of exogenous substrate N present as different N fractions in soil and exogenous substrate N uptake in plant parts was calculated using the following formula [30].

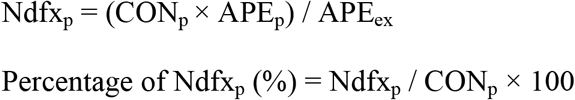

where Ndfx is N derived from urea or straw, CON is the concentration of N (mg N kg^−1^), the subscript P is the soil or plant N pool, APE is ^15^N atom percent excess, calculated by subtracting the ^15^N natural abundance from exogenous substrate N (0.365%) and ^15^N abundance of the control treatment from the applied exogenous substrate N treatments. The subscript ex is the exogenous substrate N, which is the applied urea N or straw N.

Here, the recovery rate in the fractions is the percentage of labeled exogenous substrate N (urea N or straw N) present in the target fraction at the end of the experiment [30].

### 2.6. Statistical analysis

Data are presented as the mean of three replicates. All the variables were analyzed using the SPSS 16.0 software package (SPSS Inc., Chicago, IL) at the 0.05 level. Student’s t-test was used at each growth stage to compare the differences in all variables of specific N pools between urea-only and urea plus straw treatments as well as between bulk soil and rhizosphere soil, and least significant difference (LSD) at the 0.05 protection level was used to assess the differences in mean values of the recovery rate among ^15^N-labeled urea, ^15^N-labeled urea plus straw and ^15^N-labeled straw plus urea treatments at maturity.

## 3. Results

### 3.1. Exogenous substrate N uptake in different plant parts

Under the equivalent N amendments the urea plus straw treatment significantly (P < 0.05) decreased exogenous substrate N uptake in shoots, grains, and roots at different growth stages except for shoots at the V8 stage compared with the urea-only treatment (Fig. 1a & b) because of the lower biomass in the urea plus straw treatment (Fig. S1). Conversely, urea plus straw treatment significantly (P < 0.05) increased the percentage of exogenous N present as N uptake in shoots or roots at the V8 stage over the urea-only treatment, and the opposite trend occurred in shoots, roots, and grains from stages R1 to R6 (Fig. 1c & d).

**Fig. 1.**
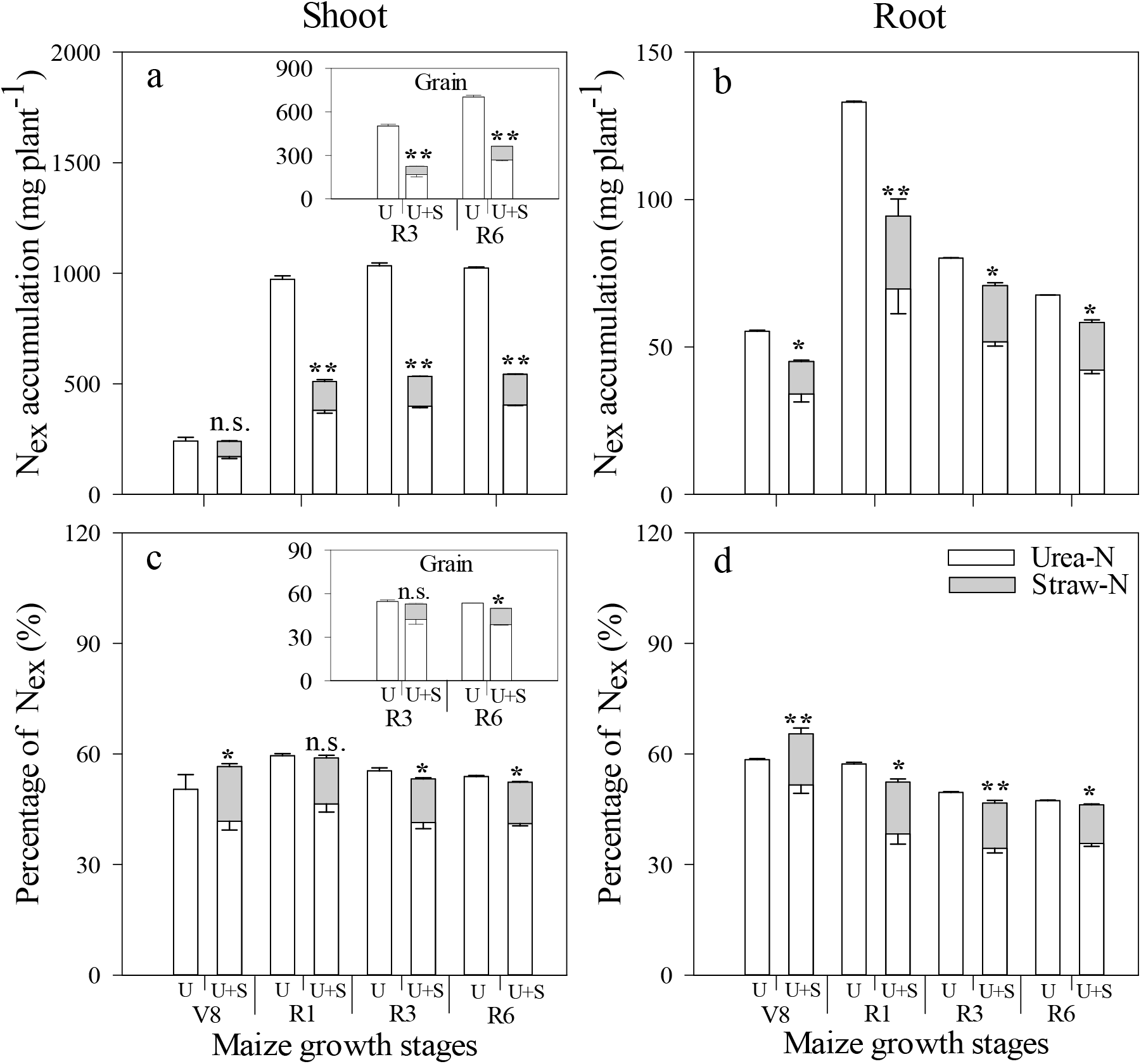
Concentration and percentage of exogenous substrate N (N_ex_) present as N taken up by maize shoots, grains, and roots under urea-only (U) and urea plus straw (U+S) treatments during maize growth stages of 8th leaf (V8), silking (R1), milk (R3), and physiological maturity (R6). Data shown are mean ± standard deviation of three replicates; * and ** denote significant differences at P < 0.05 and P < 0.01, respectively; n.s., not significant (P > 0.05). Shoot N includes leaf, stem, cob, husk, and grain fractions (all whole-plant fractions except roots)

### 3.2. Exogenous substrate N distribution in different soil N pools

Compared with the urea-only treatment, urea plus straw treatment significantly (P < 0.05) increased the concentration of exogenous substrate N present as TN in the bulk and rhizosphere soils from stages R1 to R6 except for the opposite trend in rhizosphere soil at the R1 stage (Fig. 2a & b). Similarly, compared to urea-only treatment, urea plus straw treatment significantly (P < 0.05) increased the percentage of exogenous substrate N present as TN in the bulk soil from stages V8 to R6 and in rhizosphere soil at R3 and R6 stages (Fig. 2c & d).

**Fig. 2.**
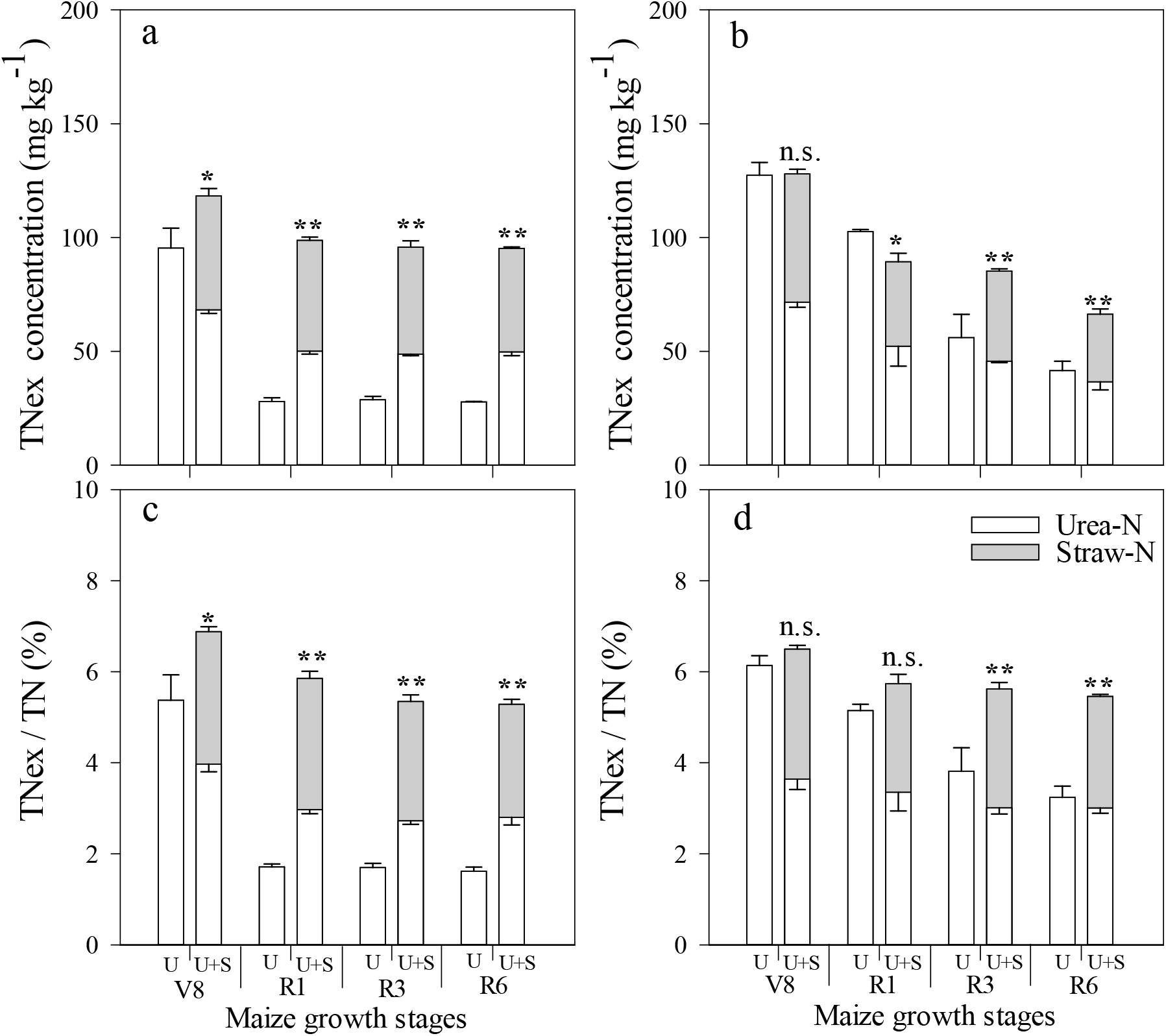
Concentration and percentage of exogenous substrate N (N_ex_) present as total N (TN) in bulk and rhizosphere soil under urea-only (U) and urea plus straw (U+S) treatments during maize growth stages of 8th leaf (V8), silking (R1), milk (R3), and physiological maturity (R6). Data shown are mean ± standard deviation of three replicates; * and ** denote significant differences at P < 0.05 and P < 0.01, respectively; n.s., not significant (P > 0.05).

Compared with urea-only treatment, urea plus straw treatment significantly (P < 0.05) decreased the concentration of exogenous substrate N present as Inorg-N in the bulk soil at the V8 stage and in rhizosphere soil at V8 and R1 stages with opposite trends in both bulk and rhizosphere soils at the R3 stage (Fig. 3a & b). However, urea plus straw treatment significantly (P < 0.05) decreased the percentage of exogenous substrate N present as Inorg-N in the bulk soil from stages V8 to R1 and in the rhizosphere soil at V8 and R1 stages, and significantly (P < 0.05) increased that in the bulk and rhizosphere soils at the R3 stage compared with the urea-only treatment (Fig. 3c & d).

**Fig. 3.**
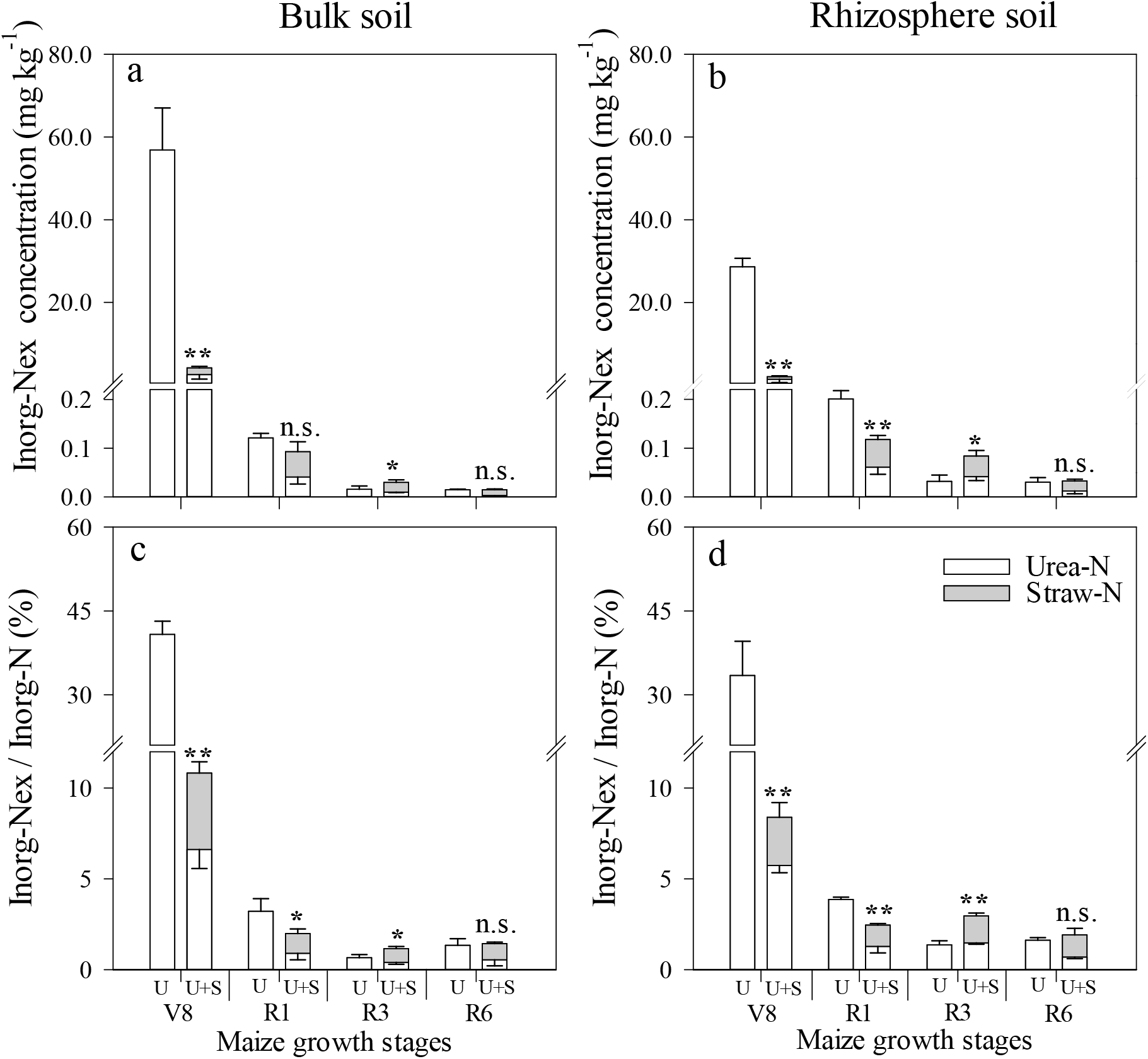
Concentration and percentage of exogenous substrate N (N_ex_) present as inorganic N (Inorg-N) in bulk and rhizosphere soil under urea-only (U) and urea plus straw (U+S) treatments during maize growth stages of 8th leaf (V8), silking (R1), milk (R3), and physiological maturity (R6). Data shown are mean ± standard deviation of three replicates; * and ** denote significant differences at P < 0.05 and P < 0.01, respectively; n.s., not significant (P > 0.05).

Compared with urea-only treatment, urea plus straw treatment significantly (P < 0.05) increased the concentration of exogenous substrate N present as DON in the bulk soil at R3 and R6 stages and in the rhizosphere soil from stages V8 to R6 except for stage R1 (P < 0.01, Fig. 4a & b). A similar trend was found in the percentage of exogenous substrate N present as DON in bulk and rhizosphere soils, but both treatments showed no significant difference in rhizosphere soil at V8 and R1 stages (Fig. 4c & d).

**Fig. 4.**
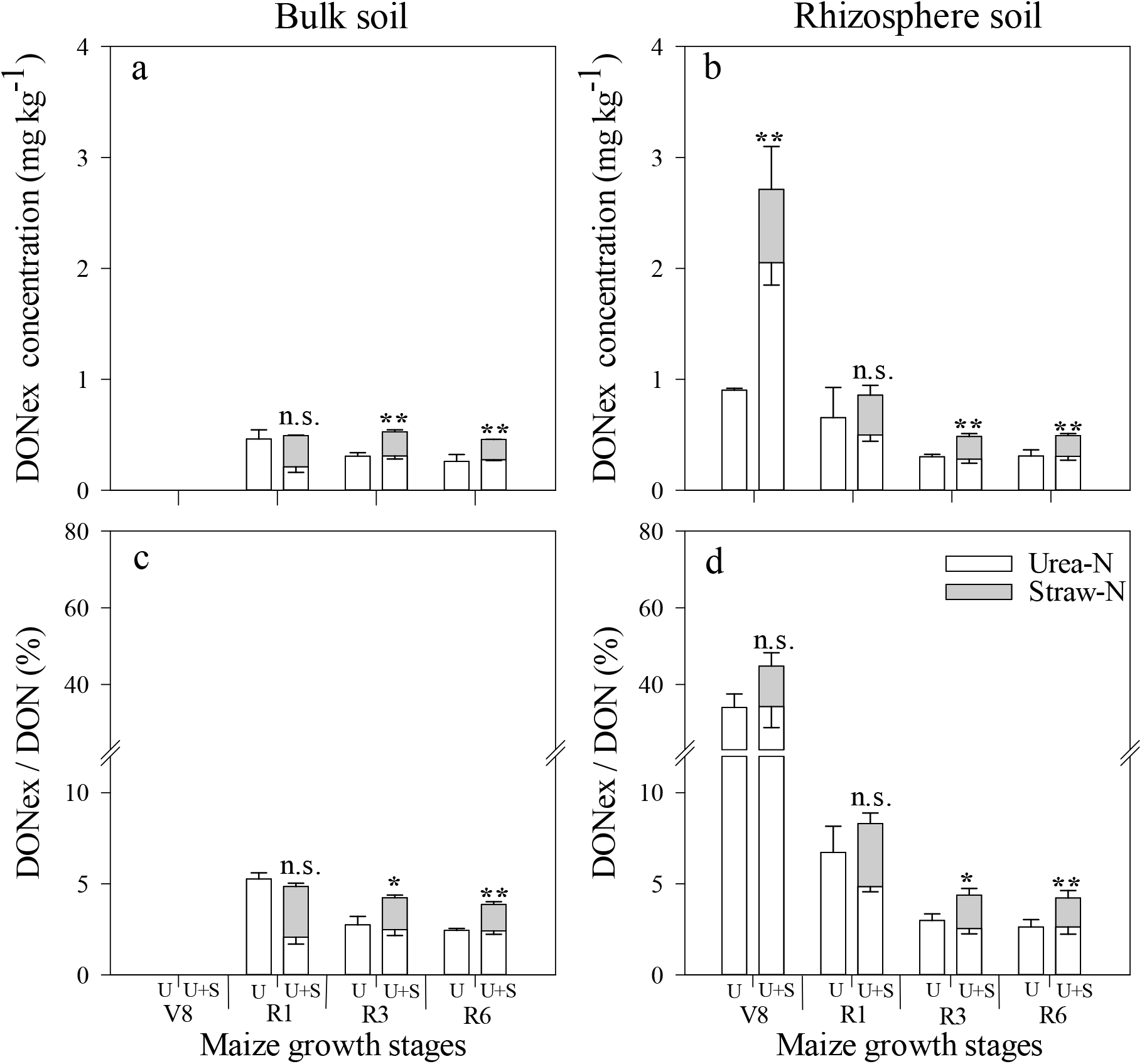
Concentration and percentage of exogenous substrate N (N_ex_) present as dissolved organic N (DON) in bulk and rhizosphere soil under urea-only (U) and urea plus straw (U+S) treatments during maize growth stages of 8th leaf (V8), silking (R1), milk (R3), and physiological maturity (R6). Data shown are mean ± standard deviation of three replicates; * and ** denote significant differences at P < 0.05 and P < 0.01, respectively; n.s., not significant (P > 0.05). Not detected in bulk soil at the V8 stage.

Urea plus straw treatment showed a significantly higher (P < 0.05) concentration of exogenous substrate N present as MBN in the bulk soil from stages R1 to R6 and that in rhizosphere soil at the R1 stage than in the urea-only treatment (Fig. 5a & b). Similarly, urea plus straw treatment showed a significantly higher (P < 0.05) percentage of exogenous substrate N present as MBN in the bulk soil from stages R1 to R6 than the urea-only treatment and the opposite trend occurred in rhizosphere soil at R6 stage (Fig. 5c & d).

**Fig. 5.**
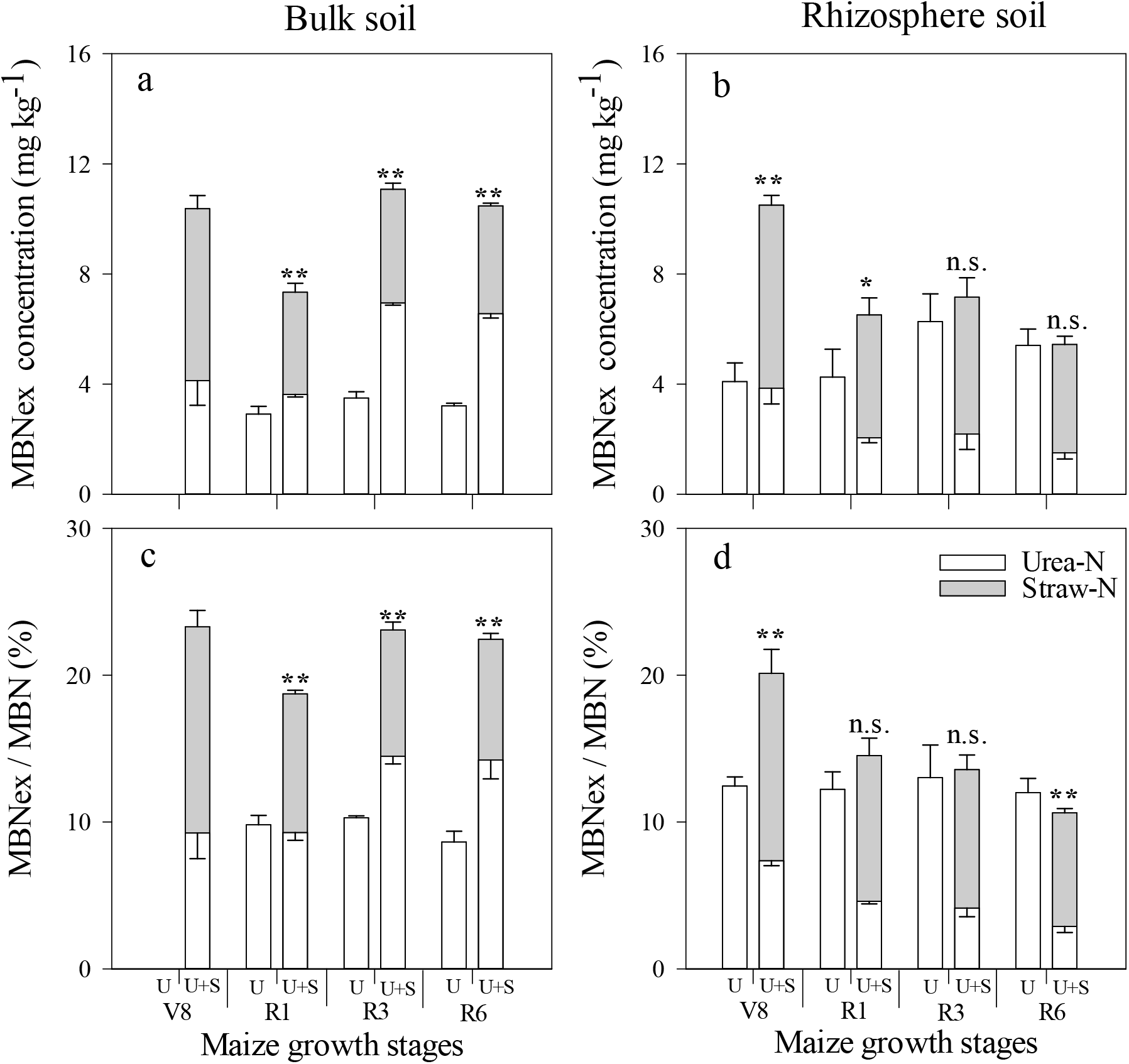
Concentration and percentage of exogenous substrate N (N_ex_) present as microbial biomass N (MBN) in bulk and rhizosphere soil under urea-only (U) and urea plus straw (U+S) treatments during maize growth stage of 8th leaf (V8), silking (R1), milk (R3), and physiological maturity (R6). Data shown are mean ± standard deviation of three replicates; * and ** denote significant differences at P < 0.05 and P < 0.01, respectively; n.s., not significant (P > 0.05).

Compared with urea-only treatment, urea plus straw treatment significantly (P < 0.01) increased the concentration of exogenous substrate N present as PON and the percentage of exogenous substrate N present as PON in the bulk soil and rhizosphere soil from stages V8 to R6 (P < 0.01, Fig. 6). The same trend in MTN was found (Fig. 7).

**Fig. 6.**
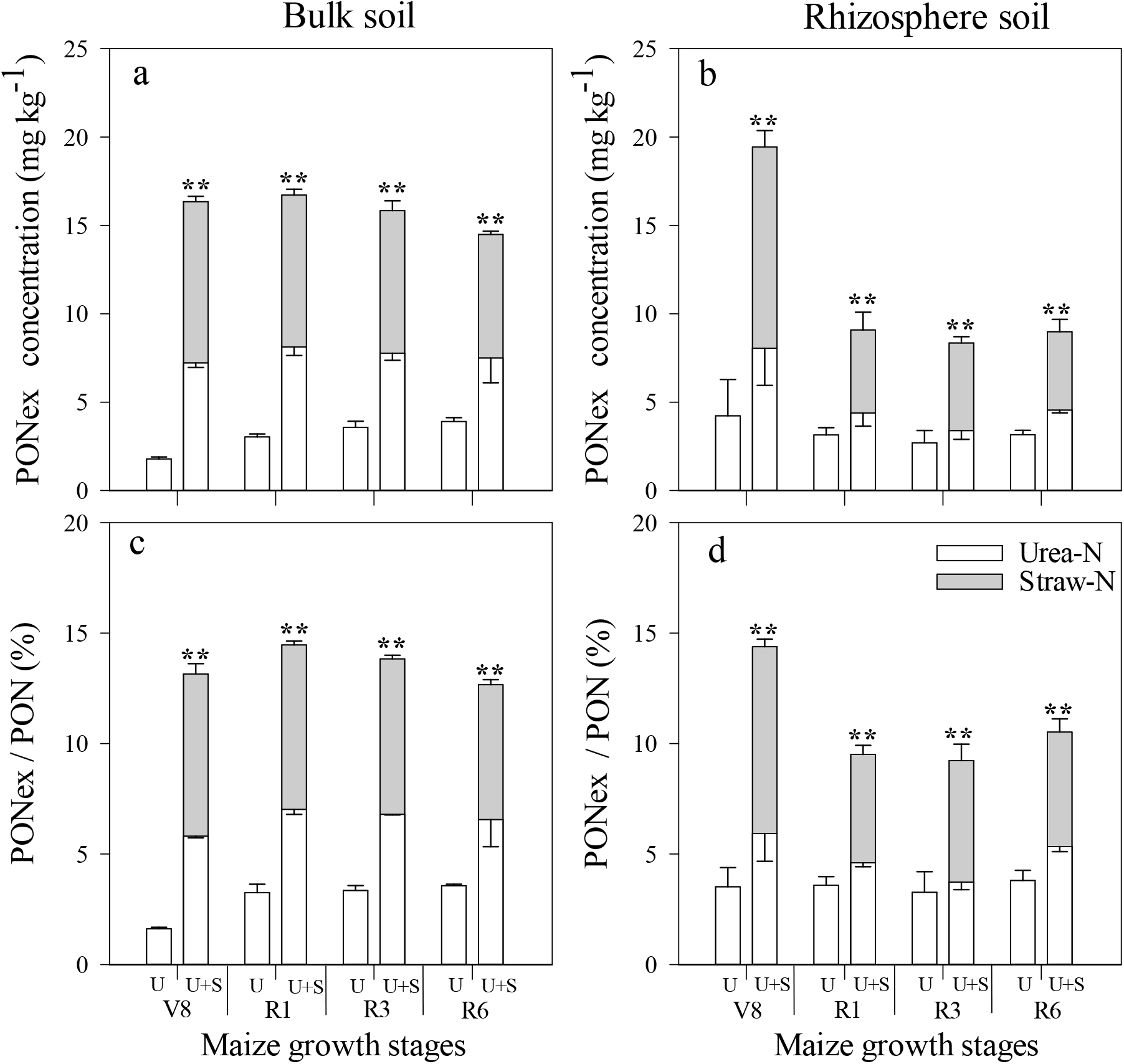
Concentration and percentage of exogenous substrate N (N_ex_) present as particulate organic N (PON) in bulk and rhizosphere soil under urea-only (U) and urea plus straw (U+S) treatments during maize growth stages of 8th leaf (V8), silking (R1), milk (R3), and physiological maturity (R6). Data shown are mean ± standard deviation of three replicates; * and ** denote significant differences at P < 0.05 and P < 0.01, respectively; n.s., not significant (P > 0.05).

**Fig. 7.**
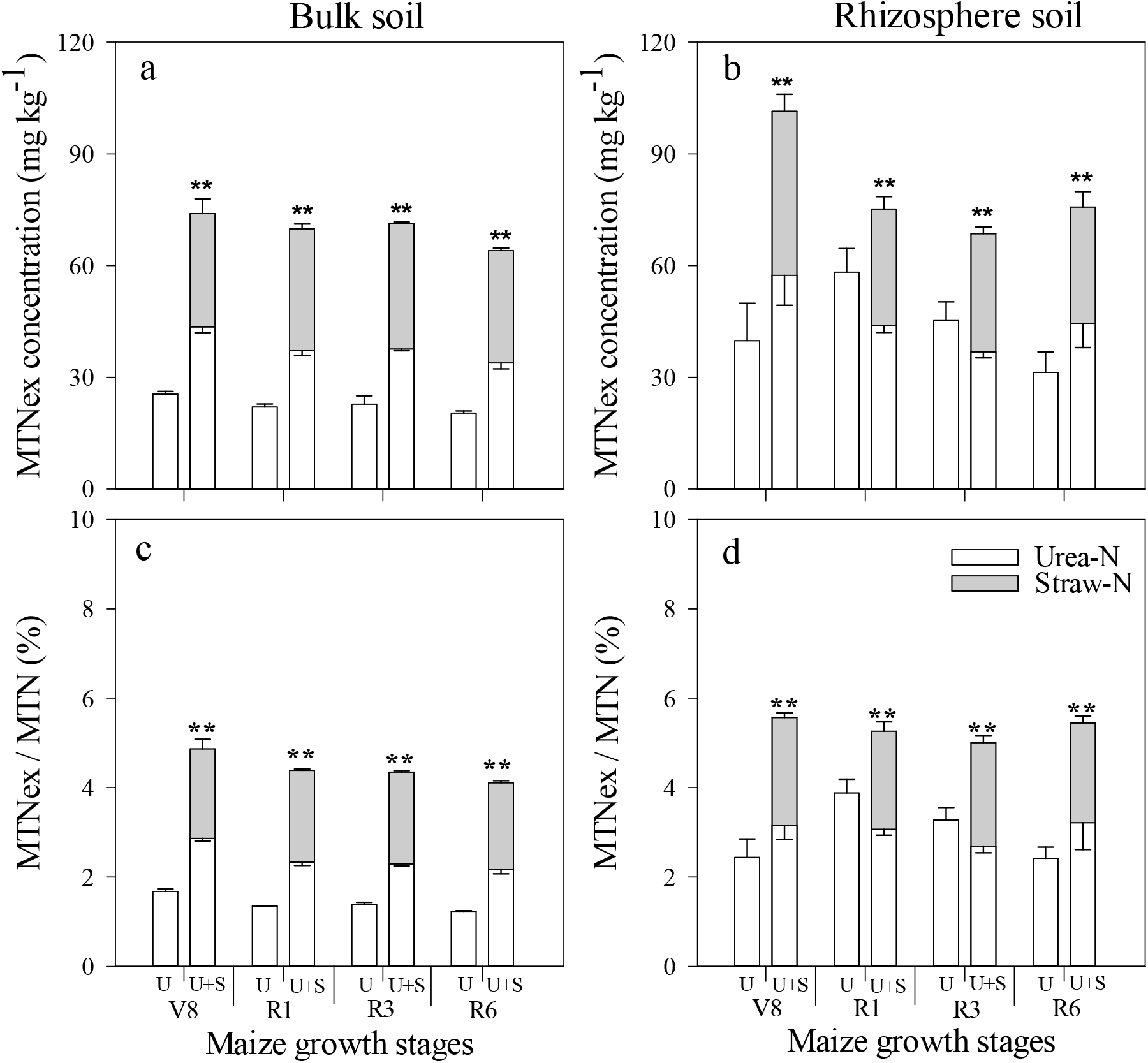
Concentration and percentage of exogenous substrate N (N_ex_) present as mineral associated total N (MTN) in bulk and rhizosphere soil under urea-only (U) and urea plus straw (U+S) treatments during maize growth stages of 8th leaf (V8), silking (R1), milk (R3), and physiological maturity (R6). Data shown are mean ± standard deviation of three replicates; * and ** denote significant differences at P < 0.05 and P < 0.01, respectively; n.s., not significant (P > 0.05).

### 3.3. Exogenous substrate N recovery in plant parts and soil N pools

As Table 2 shown, the significant (P < 0.05) sequence of exogenous substrate N recovery in plant parts was: ^15^NU > ^15^NU + S > ^15^NS + U except for exogenous substrate N recovery in the roots between ^15^NU and ^15^NU + S. The recovery rates of labeled exogenous substrate N in the whole plant, i.e., the N use efficiencies of shoots plus roots, were 71.5%, 54.9%, and 26.0% in ^15^NU, ^15^NU + S, and ^15^NS + U treatments, respectively. The N loss of urea N was 10.0% and 4.6% in ^15^NU and ^15^NU + S treatments, and no N loss was found in ^15^NS + U treatment. The significant (P < 0.05) order of exogenous substrate N recovery in soil N pools TN, PON, and MTN was: ^15^NS + U > ^15^NU + S > ^15^NU; and that in DON, MBN, and Inorg-N was: ^15^NU + S, ^15^NS + U > ^15^NU, ^15^NU + S > ^15^NS + U > ^15^NU, and ^15^NU + S > ^15^NS + U, ^15^NU, respectively.

**Table 2.** Recovery rate of labeled fertilizer N (N_L_) in different maize parts (shoots, grains, and roots) and different soil N pools (TN, Inorg-N, DON, MBN, PON, and MTN) at maturity. Unit: %.

| Treatment | Plant parts |  |  | Whole plant | Soil pools |  |  |  |  | Soil total N | N loss |
| --- | --- | --- | --- | --- | --- | --- | --- | --- | --- | --- | --- |
| | Root N / $N_L$ | Shoot N / $N_L$ | Grain N / $N_L$ | | Inorg-N / $N_L (\times 10^{-2})$ | DON / $N_L (\times 10^{-2})$ | MBN / $N_L$ | PON / $N_L$ | MTN / $N_L$ | | |
| $^{15}\text{NU}$ | 4.5 ± 0.1a | 67.0 ± 1.1a | 46.9 ± 0.8a | 71.5 ± 2.3a | 1.2 ± 0.07ab | 20.0 ± 0.4b | 2.1 ± 0.05c | 2.6 ± 0.2c | 13.6 ± 0.4c | 18.5 ± 0.8c | 10.0 ± 1.1a |
| $^{15}\text{NU} + \text{S}$ | 5.3 ± 0.2a | 49.6 ± 2.1b | 32.7 ± 1.7b | 54.9 ± 1.7b | 0.8 ± 0.06b | 33.5 ± 2.8a | 7.5 ± 0.8a | 8.3 ± 1.6b | 37.7 ± 1.8b | 40.5 ± 1.1b | 4.6 ± 0.5b |
| $^{15}\text{NS} + \text{U}$ | 2.7 ± 0.1b | 23.3 ± 1.6c | 15.7 ± 1.3c | 26.0 ± 1.1c | 1.9 ± 0.04a | 32.3 ± 3.0a | 6.5 ± 0.2b | 11.7 ± 0.3a | 50.2 ± 1.2a | 74.9 ± 2.3a | 0 |

Values are mean ± standard deviation of three replicates. Shoot N includes leaf, stem, cob, husk, and grain fractions
TN, soil total N; Inorg-N, inorganic N; DON, dissolved organic N; MBN, soil microbial biomass N; PON, soil particulate organic N; MTN, mineral associate total N; $^{15}\text{NU}$ , labeled urea-only; $^{15}\text{NU} + \text{S}$ , labeled urea + straw; $^{15}\text{NS} + \text{U}$ , labeled straw + urea; $N_L$ , labeled fertilizer N.

## 4. Discussion

### With or without straw amendment

The combination of straw and chemical fertilizer increases N recovery and decreases the loss of chemical fertilizer N [31, 32]. At the equivalent N rate in the ^15^NU-S treatment (Table 2), the N recovery (the sum of root, shoot and soil) was significantly higher at 5.4% than in the urea-only treatment at the R6 stage. However, the significantly (P < 0.05) higher N use efficiency of maize in the urea-only treatment than the other two treatments is attributed to the small amount of available N in the urea plus straw treatments in our pot experiment (Fig. 3). Some studies show that crop biomass increased with increasing crop N uptake up to the optimum N rate [33, 34]; in the urea plus straw treatment, the shortage of N supply resulted in a significantly (P < 0.05) lower shoot biomass from stages R1 to R6 and grain biomass from stages R3 to R6 compared with the urea-only treatment (Fig. S1). Conversely, the lower N supply stimulated an increase in maize root biomass to meet maize N demand [35, 36], this resulted in the significant (P < 0.05) increase of maize roots biomass at stages R1 and R3 in the urea plus straw treatment than the urea-only treatment (Fig. S1). In our study, 26.0% straw N recovery was shown in maize at the maturity, moreover, a substantial amount of recalcitrant N in the straw [37, 38] under this equivalent N amendment experiment was mainly responsible for the significantly (P < 0.05) lower 39.5 - 47.8% exogenous substrate N uptake in the shoots from the R1 stage, 11.6 - 29.1% of exogenous substrate N uptake in the roots from the V8 stage in the urea plus straw treatment compared with the urea-only treatment (Fig. 1a & b), and the lower straw N recovery in different maize parts in the ^15^NS + U treatment than the other two treatments (Table 2).

Straw has a high C/N ratio and it provides C sources for microbial N turnover and promotes the immobilization of Inorg-N. Moreover, the recalcitrant N in straw is an important stable N source in soils [39, 40], Thus, the urea plus straw treatment significantly (P < 0.05) increased the percentage of exogenous substrate N present as TN by 0.3-2.4 times in bulk soil from stages V8 to R6 and 0.5 - 0.7 times at stages R3 and R6 in rhizosphere soil compared to the urea-only treatment (Fig. 2). The same explanation accounts for the significantly (P < 0.05) higher concentration and percentage of exogenous substrate N present as DON, MBN, PON, and MTN in the bulk soil and rhizosphere soil under the urea plus straw treatment than in the urea-only treatment at different growth stages (Figs. 4–7), as well as the significant (P < 0.05) difference in N recovery in different soil N pools among ^15^NU, ^15^NU + S, and ^15^NS + U treatments (Table 2). Conversely, the characteristics of the organic N in straw under equivalent N amendment conditions was responsible for the significantly (P < 0.05) higher concentration and percentage of exogenous substrate N present as Inorg-N in the urea-only treatment compared with the urea plus straw treatment at stages V8 and R1 in the bulk soil and rhizosphere soil (Fig. 3). However, at the R3 stage in the bulk soil and the rhizosphere soil the significantly (P < 0.05) higher concentration and percentage of exogenous substrate N present as Inorg-N in the urea plus straw treatment than that in the urea-only can be attributed to the re-mineralization of the labile organic N by the mineralization-immobilization pathway and the decomposition of DON, MBN and dead roots.

### Rhizosphere and bulk soils

The rhizosphere is a zone of high nutrients and energy for soil, microorganisms, and roots [41, 42]. On the one hand, maize growth assimilated a substantial amount of N from the soil via the roots, promoting the flow of urea N from the bulk soil to the rhizosphere soil [43, 44]. On the other hand, the rhizosphere has a large amount of photosynthetically deposited C which promoted Inorg-N assimilation by microorganisms [45, 46]. With these combined effect, the urea-only treatment significantly (P < 0.05) increased the concentration and percentage of exogenous substrate N present as TN by 1.0 - 2.0 and 0.5 - 2.7 times as well as that present as Inorg-N by 0.2 - 1.1 and 0.4 - 1.0 times in rhizosphere soil in comparison with the bulk soil from stages R1 to R6 (Figs. 2 & 3, Table S1). In parallel, during rapid crop growth (V8), both inorganic N and low-molecular-weight compounds can be assimilated by crops [47–49], and the shortage of available N resulting in competition for N between maize and microorganisms may be responsible for the disappearance of exogenous substrate N in the DON in both treatments and in MBN in the urea-only treatment in the bulk soil (Figs. 5–6a & b) [50, 51].

As maize growth proceeded, a shortage of available N in the soil enhanced the competition for N between plants and microorganisms, further resulting in significantly (P < 0.05) less urea N present as MBN in rhizosphere soil than that in the bulk soil under the urea plus straw treatment, as shown by the concentration of urea N present as MBN in rhizosphere soil being significantly (P < 0.05) lower 6.7 - 77.0% than that in the bulk soil from stages V8 to R6 (Fig. 5, Table S1). Moreover, N insufficiency can stimulate the decomposition of native soil N to meet the demand of the maize and microorganisms for N [52–54]. These conclusions are supported by the observed decrease in the concentration and percentage of exogenous substrate N present as PON and the increase in the percentage of straw N present as Inorg-N in rhizosphere from stages R1 to R6 in comparison with the bulk soil (Fig. 6b & d, Table S1).

### Maize growth stages

At the V8 stage the significantly (P < 0.05) higher 18.6% of exogenous substrate N uptake in roots in the urea-only treatment than the urea plus straw treatment, and the non-significant difference in exogenous substrate N uptake in the shoots in both treatments may be attributed to the rapid growth of maize accompanied by the N transfer from roots to shoots [4] and the higher available N supply in the urea-only treatment than the urea plus straw treatment (Fig. 3). As plant growth proceeded, shoot N uptake from exogenous substrate showed a stable trend from stage R3 to R6 while root N uptake from exogenous substrate showed decrease from stage R1 because of some of the roots senescing and further leading to a decrease in root biomass after stage R1 (Fig. S1) [55, 56]. From stage R1 a slight decrease in the percentage of exogenous substrate N present in the shoots under either treatment (Fig. 1c) may have been due to the declining concentration of exogenous substrate N (Fig. 2a) and more native soil N mineralization to meet crop N demand as shown by the low Inorg-N concentration in Fig. 3. In addition, this is the contribution of native soil N for crop N uptake. Here, 40.0 - 52.5% of the percentage of native soil N uptake present as shoot N uptake in both N treatments during maize growth calculated according to Fig. 1c.

## Conclusions

The combination of urea and straw affected the distribution of exogenous substrate N in soil-crop system. Compared to the urea-only treatment at an equivalent N application rate the urea-N recovery increased by 5.4% in the urea plus straw treatment and straw recalcitrant N in the urea plus straw treatment limited exogenous substrate N uptake in maize despite the 26.0% straw N recovery in maize at the maturity (Fig. 1, Table 2). Moreover, straw labile C in the urea plus straw treatment promoted the immobilization of urea N and the contribution of exogenous substrate N to different soil N pools as shown by the significantly (P < 0.05) higher percentages of straw N and urea N in DON, MBN, PON, or MTN during the maize life cycle in most cases (Figs. 3–7). Rhizosphere soil further altered the contribution of exogenous substrate N to different N pools, e.g. rhizosphere soil showed a higher percentage of straw N and urea N present as Inorg-N and MTN in each treatment compared with the bulk soil from R1 to R6 stages (Fig.3 c, d; Fig. 7c, d). Overall, straw N availability needs to be considered after the combined application of chemical fertilizer and straw in agroecosystems, and the increase in straw N availability should be take into account the interactions between fertilization practices and the rhizosphere.

## Acknowledgements

This study was supported by the National Natural Science Foundation of China (41101277), the National Key Research and Development Plan (2018YFD0200804, 2016YFD0200101, 2018YFD0201001), and the Fundamental Research Funds for Central Non-Profit Scientific Institutions (1610132019014).

